# Isolation of an antimicrobial resistant, biofilm forming, *Klebsiella grimontii* isolate from a re-usable water bottle

**DOI:** 10.1101/724971

**Authors:** Alasdair TM Hubbard, Jesús Reiné, Enas Newire, Elli Wright, Emma A Murphy, William Hutton, Adam P Roberts

## Abstract

A re-usable water bottle was swabbed as part of the citizen science project Swab and Send and a *Klebsiella grimontii* isolate was recovered on chromogenic agar and designated SS141. Whole genome sequencing of SS141 showed it has the potential to be a human pathogen as it contains the biosynthetic gene cluster for the potent cytotoxin, kleboxymycin, and genes for other virulence factors. The genome also contains *bla*OXY-6-4 and *fosA* which is likely to explain the observed resistance to ampicillin, amoxicillin and fosfomycin. We have also shown that SS141 is a potent biofilm former, providing a reasonable explanation for its ability to colonise a re-usable water bottle. With the increasing use of re-usable water bottles as an alternative to disposables, and a strong forecast for growth in this industry over the next decade, this study highlights the need for cleanliness comparable to other re-usable culinary items.

## Introduction

As the global antimicrobial resistance (AMR) crisis continues and exacerbates, the scientific community are engaged in not only trying to discover novel antimicrobials to treat antimicrobial resistant infection [1, 2], but also understand the drivers of resistance and potential reservoirs of resistant pathogens [3–5].

*Enterobacteriaceae* are of increasing concern, so much so they have been deemed “critical” on the WHO global priority list of antimicrobial resistant pathogens. The pathogen *Klebsiella grimontii* is of particular importance as, although it can be part of the normal microflora of the gastrointestinal tract, it has been found to be an etiological agent of antimicrobial-associated haemorrhagic colitis [6]. Until recently, *K. grimontii* was thought to be a phylogroup of *Klebsiella oxytoca*, however it was found to be a distinct pathogenic species of *Klebsiella* and renamed *K. grimontii* and characterised in part by the presence of the chromosomally located beta-lactamase gene, *bla*OXY-6 [6, 7].

As part of the citizen science project Swab and Send [8], a swab was taken from a re-usable water bottle from the offices of the Telegraph Newspaper [9]. When this swab was plated out on chromogenic agar, there was an abundance of growth from presumptive *Enterobacteriaceae*. Pure culture and sequencing showed it was likely to be an antibiotic resistant, pathogenic strain of *K. grimontii*, which is able to form biofilms readily on abiotic surfaces.

## Materials and Methods

### Media and Antimicrobials

Ampicillin (AMP), amoxicillin (AMX), amoxicillin-clavulanic acid (AMC), ceftriaxone (CEF), olaquindox (OLA) and fosfomycin (FOS) were prepared in molecular grade water, ciprofloxacin (CIP) was prepared in 0.1N hydrochloric acid solution, tetracycline was prepared in 50% ethanol (all Sigma, UK) and chloramphenicol (CHL) (Sigma, UK) was prepared in ethanol (VWR, USA) all to a stock concentration of 1 mg/ml.

Growth of SS141 from −80°C stocks was on LB (Lennox) agar at 37°C for 18 hours and sub-cultured into liquid culture using either LB broth (Lennox) or cation adjusted Mueller Hinton Broth (all Sigma, UK) at 37°C for 18 hours at 200 rpm, unless stated.

### Isolation of SS141

A sample was taken from a re-usable water bottle using a Transwab^®^ Amies Charcoal Swab (MWE Medical Wire & Equipment, UK) and kept at ambient temperature. The swab was used to directly inoculate CHROMagar™ Orientation (CHROMagar, France) chromogenic agar plate and incubated at 37°C for 18 hours. A single colony that was metallic green/blue in appearance was chosen and pure-cultured onto CHROMagar™ Orientation agar. The pure isolate was then kept in stocks at −80°C in 40% glycerol (Sigma, UK).

### Sequencing and Bioinformatics

Sequencing of isolate SS141, subsequent read trimming and taxonomic distribution was performed by MicrobesNG (MicrobesNG, UK). The whole genome sequence of SS141 was assembled with SPAdes (version 3.12.0) [10] and the quality statistics of each assembly determined using QUAST (version 4.6.3) [11] and the subsequent assembled contigs were annotated using Prokka (version 1.12) [12]. Plasmids were assembled with PlasmidSPAdes (version 3.12.0) [13] using the trimmed, paired-end reads. Complete circular and linear plasmid sequences were identified by Bandage analysis through visualising the assembly graphs of the de novo genome sequences using (.FASTG) files [14]. Each plasmid contig composition was further assessed with reference to the genome sequences (.Paths) text file. The plasmid open reading frames (ORFs) were annotated using PlasmidFinder database, SnapGene software (version 3.3.4) and BacMet database (version 2.0) [15–17].

Using the assembled contigs, it was initially determined whether the isolate was pathogenic using PathogenFinder (version 1.1) [18] and compared to other strains of *K. grimontii*, *K. oxytoca* and *Klesbiella pneumoniae* by calculating the Average Nucleotide Identity (ANI) using OrthoANI (version 0.93.1) [19]. Acquired antimicrobial resistance genes located in the whole genome sequence or plasmids were searched for using RESfinder (version 3.1) [17] with an identification threshold of 60% and a minimum length of 60%. Finally, biosynthetic gene clusters (BGC) in the whole genome sequence were searched for using antiSMASH (version 4.0) [20] and the kleboxymycin biosynthetic gene cluster characterised using SnapGene software (version 3.3.4).

### Biofilm assay

Liquid culture diluted 1 in 1000 in M9 (50% (v/v) M9 minimal salts (2x) (Gibco, ThermoFisher Scientific, USA), 0.4% D-glucose, 4mM magnesium sulphate (both Sigma, UK) and 0.05 mM calcium chloride (Millipore, USA)). Three 100 μl technical replicates of the diluted culture were added to a 96 well microtitre plate alongside three technical replicates of M9 only and three empty wells and incubated statically at 37°C for 24 hours. Following incubation, all culture or media was removed, and the wells washed 4-5 times with 150 μl phosphate buffered solution (PBS, pH 7.2, Gibco, ThermoFisher Scientific, USA) and then left to dry upside down for 10 minutes. The wells were then stained with 125 μl 0.1% Gram’s crystal violet solution (Sigma, UK) for 15 minutes at room temperature, then washed 4-5 times with 150 μl PBS and left to dry upside down for between 60-90 minutes. The remaining stain was dissolved with 125 μl 30% acetic acid (Sigma, UK) incubated at room temperature for 15 minutes. Finally, the acetic acid solution was transferred to a fresh 96 well microtitre plate and measured at an optical density of 550nm, with 30% acetic acid used as a blank and the stained empty wells as background.

### Minimum Inhibitory Concentrations

Minimum inhibitory concentration (MIC) of AMP, AMX, AMC, CEF, CIP, OLA, CHL and FOF to SS141 were carried out using cation adjusted Mueller Hinton Broth following the CLSI Guidelines for Antimicrobial Susceptibility Testing using the Broth Microdilution Methods.

## Results and Discussion

The swab taken from a re-usable water bottle was initially plated out on to the chromogenic CHROMagar™ Orientation agar routinely used for identification of urinary tract pathogens [21] which identified a significant amount of presumptive *Enterobacteriaceae* present on the swab. Due to the metallic blue/green pigmentation and morphology of the colonies on the chromogenic agar it was determined that these colonies were probable *Citrobacter*, *Enterobacter* or *Klebsiella*, which are often pathogenic. We therefore sub-cultured until it was a pure isolate, denoted as isolate SS141. Following whole genome sequencing, the isolate was identified as *K. grimontii*. SS141 contains three plasmids; two low copy number plasmids IncFIIK (90X coverage depth) and IncFIA(HI1) (40X coverage depth) and a small plasmid with extremely high copy number (30,000X coverage depth). Antimicrobial resistance genes were not found to be present on any of these plasmids, however, heavy metal resistance genes (*arsR*, *silE*, *cusA*, *PcoR* and *CopA/B/C/D* conferring resistance to arsenic, silver and copper, respectively) were found to be present on the IncFIA(HI1) and the small plasmid. While *K. grimontii* can be asymptomatically carried by humans, it has also been associated with the cause of a number of infections including antibiotic-associated haemorrhagic colitis and bacteraemia [6]. We were able to identity, using PathogenFinder, that the SS141 isolate is likely to be a human pathogen with a probability score of 0.842. Subsequently, we compared the SS141 genome to a clinical strain of isolated from a patients sputum *K. grimontii* WCHKG020121 [7] and the reference strain *K. grimontii* 06D021 [6], three published genomes of pathogenic strains of *K. oxytoca*; *K. oxytoca* 09-7231-1 isolated from a mouse tumour abscess [22], *K. oxytoca* E718* a clinical isolate from Taiwan [23] and *K. oxytoca* JKo3 a clinical isolate from a strain collection in Japan [24], as well as two clinical isolates of *K. pneumoniae* (KPL0.1 and KPL0.2) [25] using ANI (Fig. 1). We found that SS141 was very closely related to the clinical isolate *K. grimontii* WCHKG020121 (99.74%) and reference strain *K. grimontii* 06D021 (99.2%), further confirming that SS141 is *K. grimontii*. SS141 was also 99.22% identical to *K. oxytoca* JKo3 suggesting that *K. oxytoca* JKo3 may have been mis-identified and is *K. grimontii* (Figure 1). All these strains had a similar likelihood of pathogenicity as determined by PathogenFinder (Figure 1).

**Figure 1:**
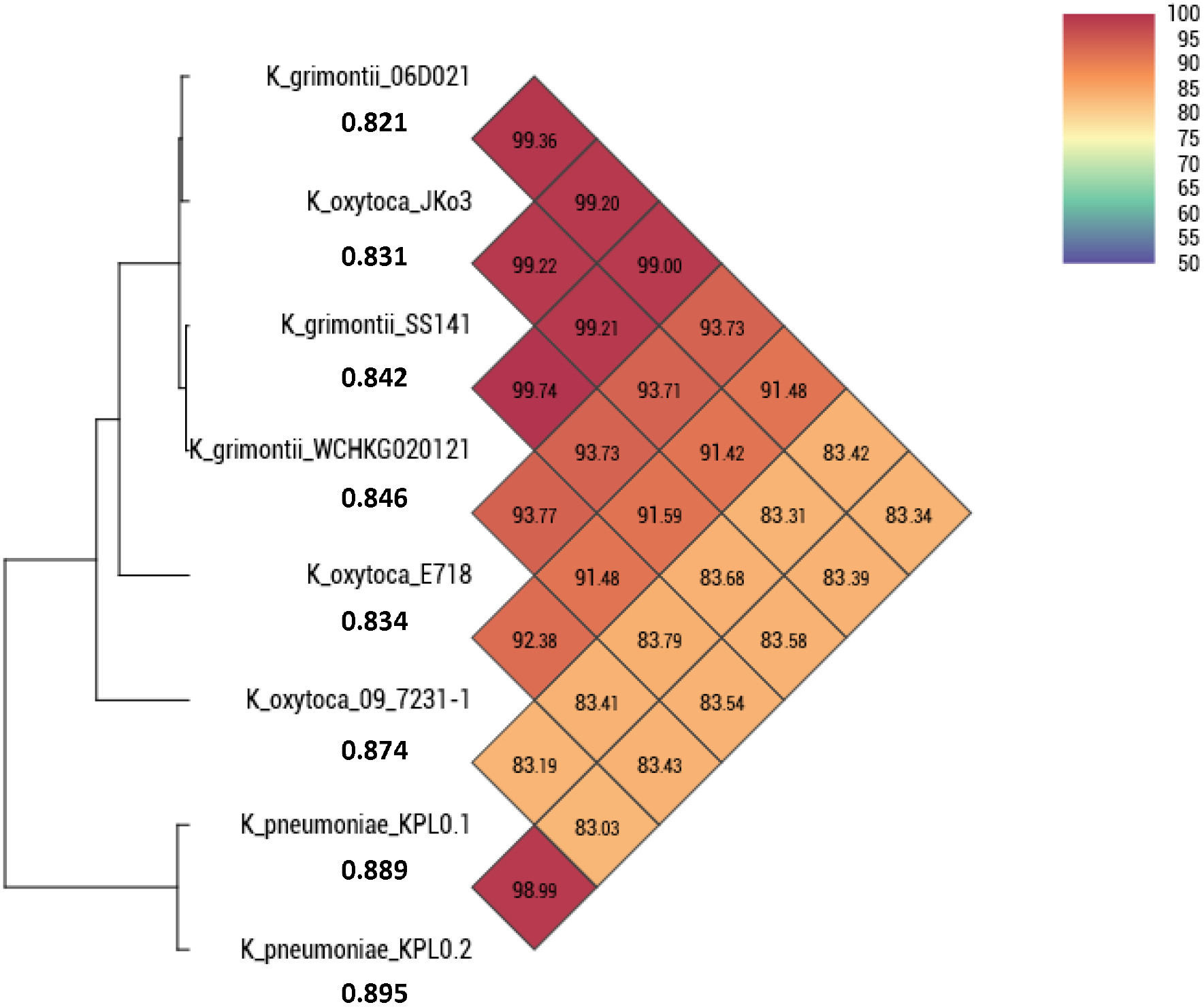
Calculation of the Average Nucleotide Identity (ANI) of *K. grimontii* SS141, *K. grimontii* WCHKG020121, *K. grimontii* 06D021, *K. oxytoca* 09-7231-1, *K. oxytoca* E718, *K. oxytoca* JKo3, *K. pneumoniae* KPL0.1 and *K. pneumoniae* KPL0.2. \**K. oxytoca* E718 was submitted as *Klebsiella michiganensis* on GenBank. Figures under the isolate name represent the probability of being a human pathogen, as determined by PathogenFinder.

Five antimicrobial resistance genes were identified to be present on the SS141 chromosome using RESfinder, *bla*OXY-6-4 which confers resistance to beta-lactam antibiotics [26] and is an indicator of *K. grimontii* [6, 7]. Both *oqxA* and *oqxB* which are involved in resistance to olaquindox [27], an antimicrobial used as a growth promoter in animals [28], and the fosfomycin resistance gene *fosA* were all also identified and were all previously found to be present on the chromosome of several other *K. grimontii* isolates [7]. Finally, *mdf(A)* was identified which has previously been shown to confer resistance to a range of different antimicrobial compounds, including chloramphenicol [29]. Resistance to the beta-lactam antimicrobials AMP and AMX were confirmed (MIC of 16 μg/ml and 32 μg/ml, respectively) however SS141 was still sensitive to both AMC (2 μg/ml) and CEF (0.0625 μg/ml) according to EUCAST clinical breakpoints (Table 1). Resistance to fosfomycin was also confirmed with a MIC of 256 μg/ml, which is well over the clinical breakpoint of >32 μg/ml. Despite the presence of *oqxAB* and *mdf(A)* genes in SS141, the isolate was sensitive to both OLA (16 μg/ml) and CHL (2 – 4 μg/ml) according to the previously determined breakpoint of >64 μg/ml [27, 30] and >8 μg/ml, respectively (Table 1). We noticed that the *oqxA* and *oqxB* genes identified by RESfinder also closely aligned with the genes encoding the efflux pumps BepF and BepE, respectively which have been shown to confer resistance ampicillin, norfloxacin, ciprofloxacin, tetracycline, and doxycycline in *Brucella suis* [31]. We tested for resistance to tetracycline however the strain was susceptible with an MIC <0.5 ug/ml (the lowest concentration tested).

**Table 1:** Minimum inhibitory concentrations of ampicillin (AMP), amoxicillin (AMX), amoxicillin-clavulanic acid (AMC), ceftriaxone (CEF), ciprofloxacin (CIP), olaquindox (OLA), chloramphenicol (CHL) and fosfomycin (FOF) to the SS141 isolate.

|  | AMP | AMX | AMC | CEF | CIP | OLA | CHL | FOF |
| --- | --- | --- | --- | --- | --- | --- | --- | --- |
| SS141 | 16 µg/ml | 32 µg/ml | 2 µg/ml | 0.03125 – 0.0625 µg/ml | 0.0078125 µg/ml | 16 µg/ml | 2 – 4 µg/ml | 256 µg/ml |

The SS141 isolate contains both *treC* and *sugE* genes, both of which have previously been shown to be involved biofilm formation in *Klebsiella pneumoniae*, with *treC* being particularly important to the colonisation of the gastrointestinal tract [32]. We have confirmed that SS141 is a potent biofilm former, producing a significant (Ordinary one-way ANOVA, uncorrected Fisher’s LSD *P* value = <0.0001) biofilm during growth in the minimal media M9 compared to *Escherichia coli* 10129 [33]; a known biofilm producer (Fig. 2). This would provide a good explanation for isolation of the Klebsiella from the swab and indicates that SS141 has the potential to persist in the environment, in this case on a reusable water bottle. Finally, the BGC for the potent cytotoxin kleboxymycin was identified using antiSMASH and found to be present within the SS141 genome with 99.53% homology with the previously characterised kleboxymycin BGC (accession number MF401554) [34]. This cytotoxin has been previously associated with the cause of antibiotic-associated haemorrhagic colitis [34], further suggesting that SS141 is a pathogenic strain of *K. grimontii*.

**Figure 2:**
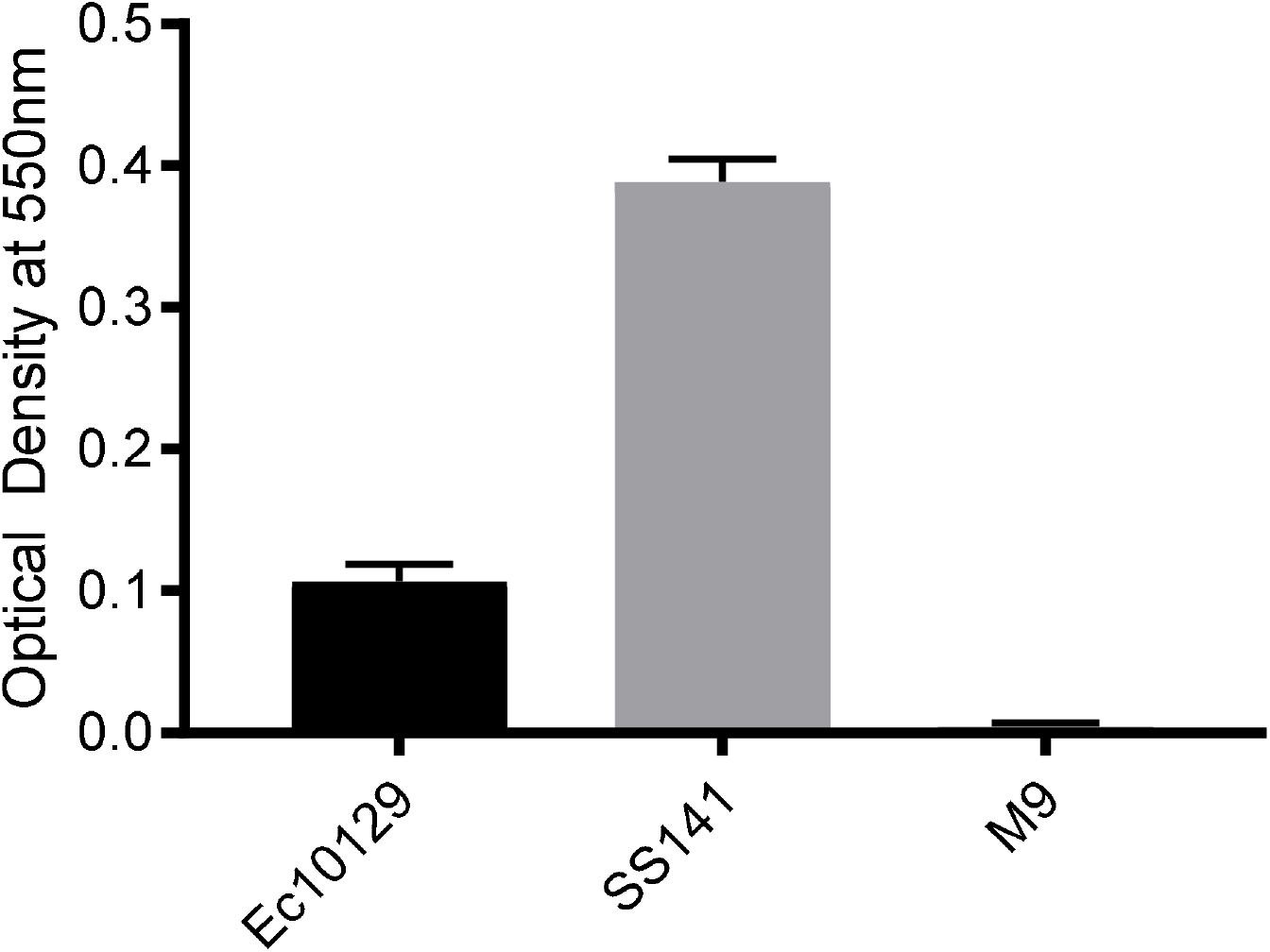
Biofilm formation of the SS141 isolate and *E. coli* 10129 in M9 media. Error bars represent the standard error of the mean.

## Conclusions

We describe the first instance of isolation of a *K. grimontii* isolate from a reusable water bottle. The isolate was found to be a potent biofilm former and carried multiple resistance genes to both metals and antibiotics. It is also likely to be pathogenic as the biosynthetic gene cluster for kleboxymycin is present within the genome. There is a strong forecast in the global market for reusable water bottles over the next decade [35] and this study highlights the seemingly overlooked aspect of cleanliness when it comes to repeated use of such household items. There is, as yet, no comprehensive scientific study of bacterial colonisation of reusable water bottles and perhaps this is needed to persuade manufacturers and users to promote and practice suitable washing regimens to keep bacterial load to a minimum.

## Accession number

This Whole Genome Shotgun project for *Klebsiella grimontii* SS141 has been deposited at GenBank under the accession RXHH00000000. The version described in this paper is version RXHH01000000.

## Acknowledgments

We would like to thank MWE, Medical Wire & Equipment, for the continued support of the Swab and Send citizen science project.

